# Electrolysis of Bacterial Biofilm

**DOI:** 10.1101/2021.05.27.445919

**Authors:** Abdollah Bazargani, Arsalan Azimi, Pouya Farhadi, Seyed Sina Naghibi Irvani, Psalm Moatari

## Abstract

Biofilms are a significant global health problem and are of most important clinical impediments of the century. Biofilms play a crucial role in antimicrobial resistance and chronic infections by acting as reservoir for microorganisms. Therefore greater understanding of biofilm and novel methods to eliminate them are needed. We considered electrolysis as a contrivance to decompose and disperse the *Pseudomonas aeruginosa biofilm* in microtiter dishes. We showed electrolysis effectively eliminated the *Pseudomonas aeruginosa biofilm with the potential to change our approach in tackling biofilms*.

## 1. Introduction

Biofilm is an assembly of microorganisms such as bacteria, fungi, viruses and/or protozoa, embedded within a matrix. Such matrix reinforces these organisms to each other and/or to a surface [1]. Many different gram-positive (e.g. *Staphylococcus* spp, *Listeria monocytogenes, Bacillus* spp, and *lactic acid bacteria* spp) and gram-negative bacterial species (e.g. *Pseudomonas aeruginosa* or *Escherichia coli*) could live in biofilm format [2]. Matrix of bacterial biofilms, a sticky hydrated gel that holds biofilm together, is a conglomeration of water and extracellular polymeric substances (EPS), a composition of proteins, polysaccharides and extracellular DNA [3]. Bacterial biofilms are capable of adhering to most of living and non-living surfaces, including living tissues, implantable medical devices, percutaneous sutures, tracheal and ventilator tubes and industrial or potable water system pipings making them prevalent in healthcare settings [4]. Compared to their planktonic counterparts, bacteria living in a biofilm obtain numerous benefits. For instance, becoming highly resistant to innate and adaptive immune systems of host, and to antibiotics and disinfectant agents [5]. Studies have shown the bacteria in a biofilm can become up to 1,000-fold more resistant to previously effective antimicrobial agents [6, 7]. As a result biofilms play a crucial role in persistence and chronicity of a wide variety of microbial infections [8] such as periodontal diseases, recurrent urinary tract infections, bacterial vaginosis, chronic sinusitis and middle-ear infections, infective endocarditis, chronic pulmonary infections in cystic fibrosis [9], and infections of permanent indwelling medical devices such as prosthetic cardiac valves and artificial joint prostheses, cardiac pacemakers, internal fixation devices, percutaneous sutures, and tracheal and ventilator tubes [8]. Implantable medical devices favor microorganism adhesion, colonization and biofilm formation. 80% of all microbial infections are biofilm related infections making them the most important clinical impediments of the century [8]. Microbiologist researchers have begun to investigate microbiologic processes from a biofilm perspective [1]. Although numerous approaches have been considered to both inhibit the formation and/or to disperse bacterial biofilms of surfaces, no available method has successfully been utilised clinically [11-13].

The matrix serves numerous functions namely: it keeps the bacteria in close proximity to each other, facilities intercellular communication, serves as nutrient source. It accounts for 90% of the biofilm’s dry mass and no wonder strategies that focus on destroying the integrity of biofilm matrix could result in new therapeutic approaches to biofilm related infections [11-14].

We have hypothesized that electrolysis could decompose matrix of biofilm. Electrolysis is a chemical reaction whereby a direct electric current (DC) passes through an ionic substance. This drives an otherwise non-spontaneous chemical reaction and separates anionic and cationic components of ionic compounds. The main parts of electrolysis are:

- **An electrolyte:** an ion-conducting liquid containing free ions that carry electric current in the electrolyte.
- **A direct current (DC) electrical supply:** providing electrical current carried by electrons in an external circuit and provides necessary energy to drive an otherwise non-spontaneous chemical reaction that separates anionic and cationic components of ionic compounds.
- **Two electrodes:** electrical conductors that do not take part in electrolytic reaction and provide physical interface between electrolyte and electrical circuit [15].

Biofilm matrix is a highly hydrated environment and water, an ion-conducting liquid, is the largest component of matrix [14]. Matrix also contains anions and cations [16]. As water in biofilm matrix together with cations and anions could act as components of electrolysis, it is hypothesized that biofilm could be subjected to electrolysis in order to disintegrate it. Here we have evaluated the efficacy of Electrolysis in decomposition of biofilm of Pseudomonas aeruginosa.

## 2. Materials and methods

In order to study possible effects of electrolysis on biofilms, and to assay an *in-vitro* biofilm, we followed “Microtiter dish biofilm formation assay”, established by “O’Toole”: [17]

1. Wild-type *Pseudomonas aeruginosa* was cultured in a rich medium.
2. 100 μL of culture was added to 9900 μL of M63 fresh minimal medium, supplemented with magnesium sulfate, arginine (Table 1).
3. Next, using a pipet, 100 μL of the dilution was transferred into 96 wells of the plate.
4. The microtiters were incubated at 37°C for 24 hrs.
5. After incubation, the cells were turned over and shook and the bacteria were dumped out of cells.
6. The plates were submerged in a small tub of water and then water was shook out. The process was performed twice.
7. Using a pipet, 125 μL of 0.1% crystal violet solution was added to each well of the microtiter plate.
8. The plate was incubated at room temperature for 15 minutes and then the plate was washed for 4 times.
9. The plate was turned upside down and dried for 24 hours.
10. A DC electrical supply that could circuit 1,000 micro-amperes of electrical current between 1.5-12 volts of potential difference was used to perform electrolysis.
11. Platinized titanium wires were used as electrodes
12. 200 μL of Sterile distilled water was added to each well.
13. Electrolysis was performed, maximally for 60 seconds, with 1000 micro-amperes of electrical current at 1.5, 3, 4.5, 6, 7.5, 9, 10.5 and 12 volts of potential difference. Each voltage electrolysis was performed twelve times and mean time for initiation of electrolysis was measured.

## 3. Results

Process of electrolysis was performed and we waited for initiation of electrolysis effects for a maximum of 60 seconds and its effects on the biofilm (appearance of gas bubbles, detachment of biofilm from surface) was evaluated with a 10X hand held magnifier. After 60 seconds, no obvious gas bubble was detected at 1.5, 3 and 4.5 volts. Although the process began after 40-50 seconds by 6 and 7.5 volts of potential difference, 30-40 seconds by 9 volts, 20-30 seconds by 10.5 volts and 10 seconds by 12 volts (Table 2).

To avoid mechanical damage to the biofilm, the electrodes were gently moved over the biofilm and process of electrolysis was continued until dispersal of whole violet rim of biofilm from the surface. The following images show steps of electrolysis of the biofilm by 12 volts of potential difference (Fig. 1-3).

**Figure 1.**
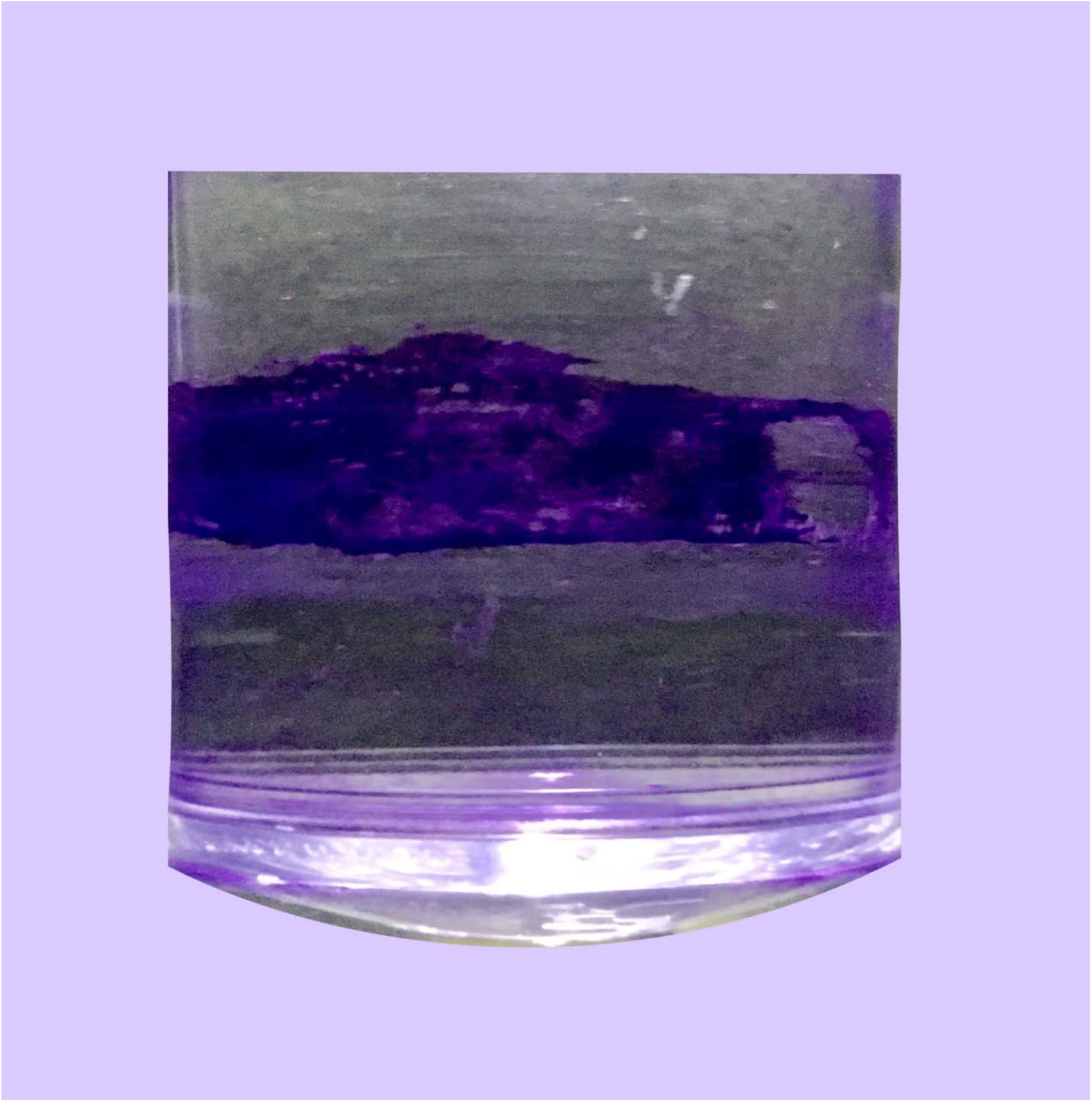
The biofilm was grown in dishes of a microtiter plate, stained with crystal violet and appeared as thin violet rims on walls of the microtiter dishes. A dish with a robust biofilm on walls was chosen and 200 μLs of sterile distilled water was added to the micro titer dish.

**Figure 2.**
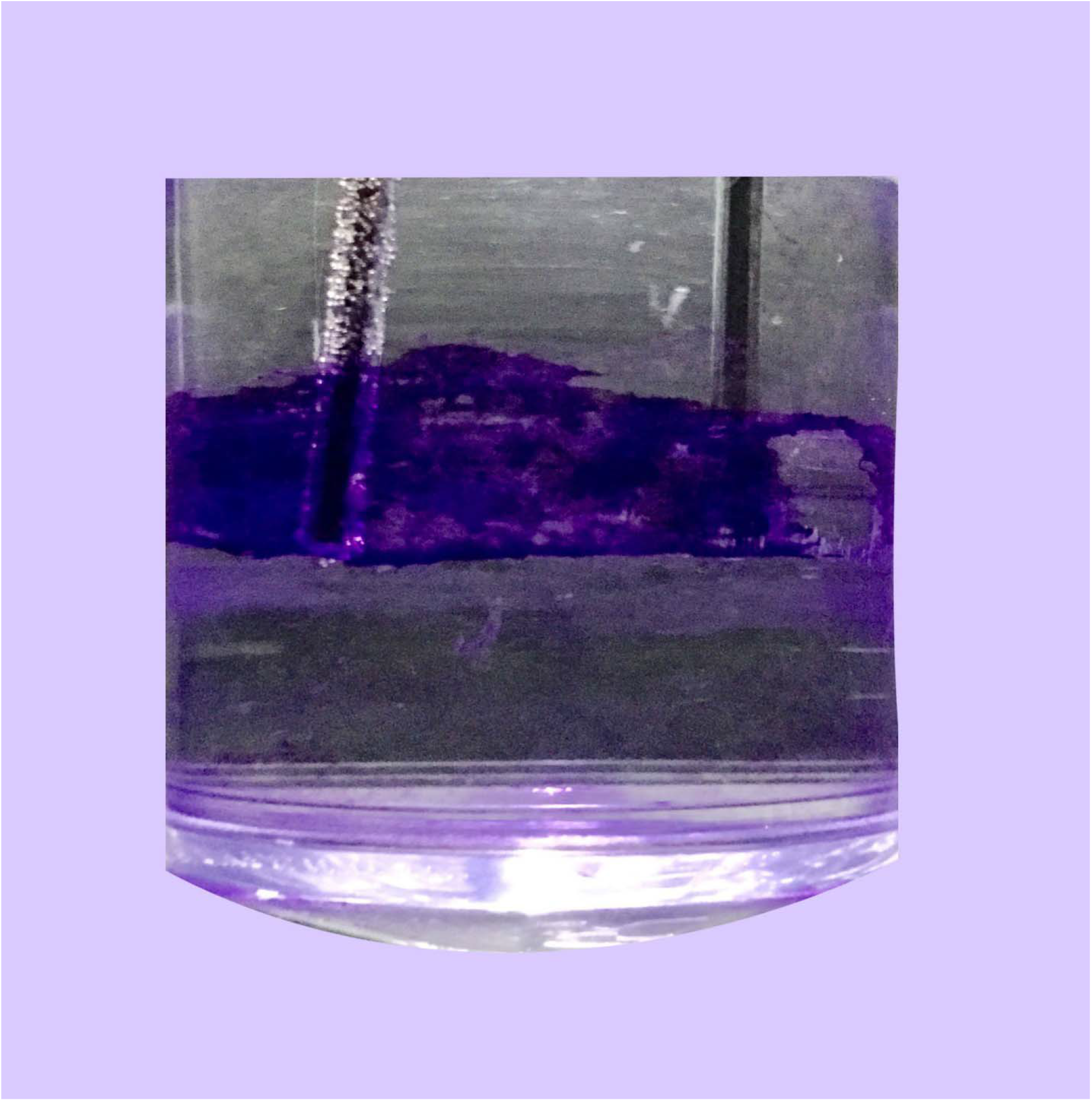
The electrodes (supplied with 12 volts, 1 ampere) were inserted on surface of the biofilm and the process of electrolysis initiated after about 10 seconds and the whole rim of biofilm was electrolyzed.

**Figure 3.**
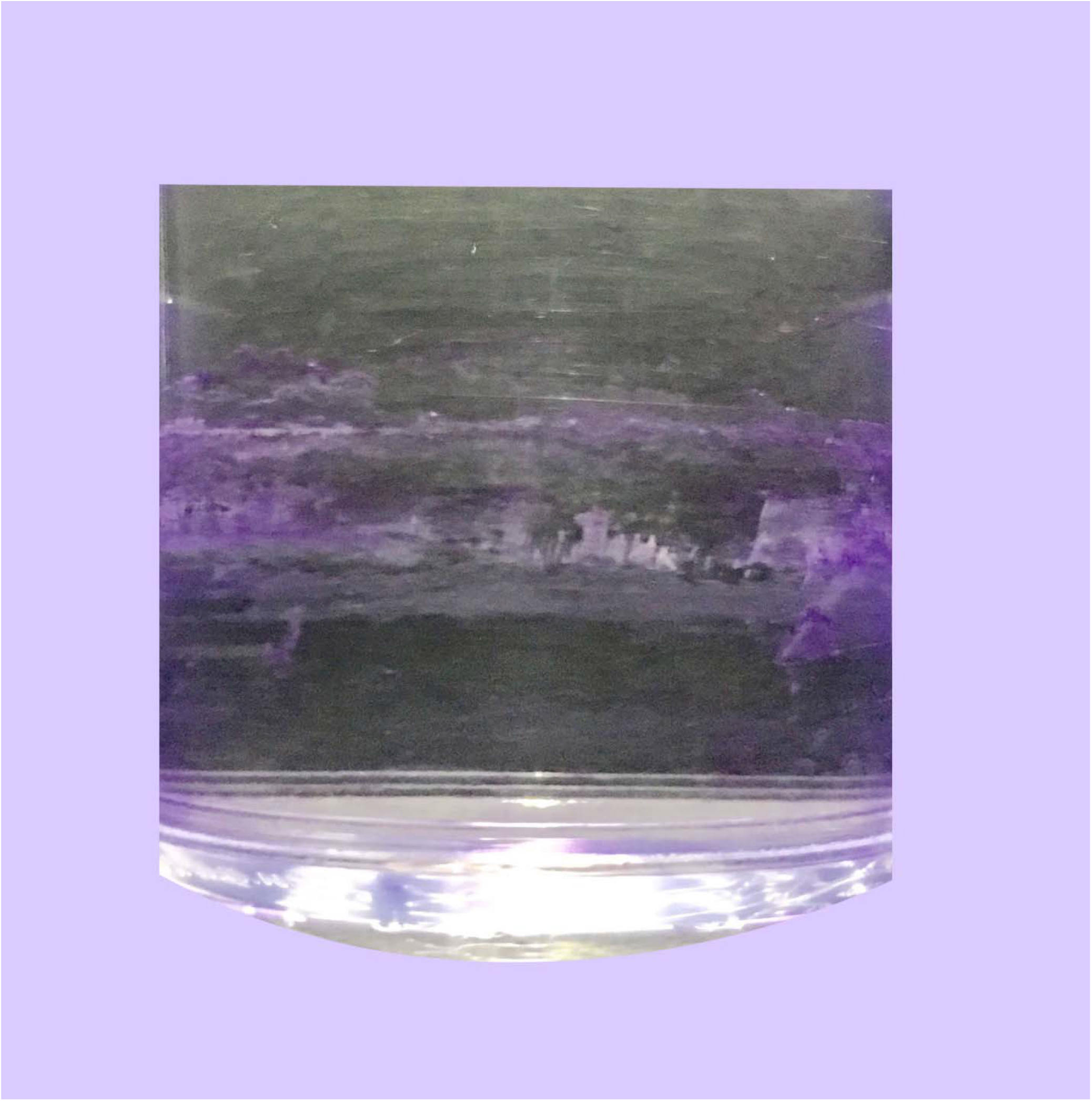
The biofilm was dispersed from the surface.

## 4. Discussion

We evaluated whether biofilm electrolysis is able to disintegrate its matrix into detached particles. We showed electrolysis can lyse and disperse biofilm and remove it from surface. As mentioned before 80% of all microbial infections are biofilm related infections [8]. Biofilm enables the bacteria to become resistant to previously effective antibiotics [1] and also to evade the host immune response [11]. “Biofilm-related chronic infections are very difficult, if not impossible, to cure wit h antibiotics.” [18]. There are three main strategies to tackle the biofilm.1) strategies that physically or chemically remove a currently formed biofilm (e.g autoclave). 2) strategies that eradicate bacteria living in a biofilm (e.g. radiation, antibiotics). To remove currently formed biofilms many different strategies have been evaluated. For example, a wide variety of antibiotics, bacteriophages [19], chemical agents [20] antimicrobial hydrogels [21] and many natural compounds known for their antimicrobial properties (such as brominated furanones [22], garlic [23], ursine triterpenes [24], corosolic acid and asiatic acid [25], ginseng [26] and 3-indolylacetonitrile [27]) have been suggested. Unfortunately, none of these methods have been approved for clinical use [13]. And third group of strategies should prevent formation of biofilm on living and/or nonliving surfaces. To prevent formation of biofilm on medical devices, surfaces of medical devices could be covered with molecules and/or polymers that impair biofilm formation. Our study is the first of its kind that evaluates the effects of electrolysis on biofilms. We showed electrolysis dispersed and lysed biofilm of *Pseudomonas aeruginosa* from walls of microtiter dishes. Rate of elimination was proportional to power of electrical supply. The method could be used in future in order to remove biofilms from different surfaces and can be used to remove biofilm from surface of indwelling medical devices, implants, chronic wounds, dental plaques, to clean water pipes of hospitals and etc. Further studies are needed in order to determine most appropriate, least harmful and most effective voltage and amperage to remove biofilms from surfaces in each application. Further studies are needed to evaluate the theatrical effect of electrolysis on the bacteria in the biofilm.

Interestingly, through stimulation of electrotaxis, electrical stimulation may promote wound healing [28]. Thus hypothetically therapeutic intervention with electrolysis may not only disperse biofilm of surface of chronic wounds, but also increase rate of wound healing and further studies could also evaluate such hypothesis.

One of limitations of our study was that electrolysis was performed with constant electrical current but different voltages, thereby further studies should be performed in order to evaluate the process with different electrical currents and at different voltages. Another limitation of our study was that before electrolysis, biofilms were stained by crystal violet which is an ionic water soluble compound that might have affected the process of electrolysis. We suggest staining biofilms after electrolysis and compare the results with a non-electrolyzed biofilm on same surface. In order to compare the results researchers could use reverse and electrical microscope and could also take samples from biofilms for bacterial culture and evaluate bactericidal properties of electrolysis.

## Conclusion

Biofilms are a significant global health problem and are of the most important clinical impediments of the century. A greater understanding of biofilm with novel methods to eliminate it are therefore needed. We showed electrolysis effectively removed biofilm from the surface. We suggest electrolysis as a new approach to tackle the biofilm. Further studies are needed to utilise this method into clinical practice.

## Acknowledgments

We would like to appreciate Zahra A. Kashkooli and Psalm Moatari for critical reading and editing the manuscript.

## Figures Legends

**Table 1** Microtiter dish biofilm formation assay

**Table 2**Time needed for initiation of electrolysis in different voltages

