## Supplementary material for "Electrolysis of Bacterial Biofilm": Tables 1,2

**Table 1**

Microtiter dish biofilm formation assay

| **Name** | **Company** | **Catalog Number** | **Comments** |
| --- | --- | --- | --- |
| 1 X M63 |  |  | First dissolve 35g K_2_HPO_4,_ 15g KH_2_PO_4_, and 10g (NH_4_)_2_SO_4_ in 1 L of water. This stock is 5X M63, Dilute 5X stock 1:5. Before adding desired components, autoclave M63, and cool, M63. |
| KH_2_PO_4_ | MERCK | 1125112A |  |
| K_2_HPO_4_ | MERCK | 1051011000 |  |
| (NH_4_)_2_SO_4_ | Sigma-Aldrich | A5132 |  |
| Magnesium sulfate | AnalaR NORMAPUR | MFCD00011110 | Prepare MgSO_4_ as a 1 M stock in water and then autoclave it. Its concentration should be 1 mM in final concentration. |
| Arginine | Sigma-Aldrich | A5131 | Arginine concentration should be 0.4% in final concentration. In order to do so Prepare it as a 20% stock in water,then filter and sterilize it. |
| Microtiter plates | BD Biosciences | 353911 | Falcon 3911, Microtest III, Flexible assay plates, 96 well, U-bottom, non-sterile, non-tissue-culture treated. |
| Microtiter plate lids | BD Biosciences | 353913 | The lids can be reused if cleansed with 95% ethanol in water. |
| Crystal violet | MERCK | 1159400100 | Its concentration should be 0.1% in water. |

**Table 2**

Time needed for initiation of electrolysis in different voltages

| Voltage _(Volts)_ | 1.5 | 3 | 4.5 | 6 | 7.5 | 9 | 10.5 | 12 |
| --- | --- | --- | --- | --- | --- | --- | --- | --- |
| Time _(seconds)_ | 0 | 0 | 0 | 40-50 | 40-50 | 30-40 | 20-30 | 10 |
